# Corticothalamic Projections Deliver Enhanced-Responses to Medial Geniculate Body as a Function of the Temporal Reliability of the Stimulus

**DOI:** 10.1101/2021.05.07.443156

**Authors:** Srinivasa P Kommajosyula, Edward L. Bartlett, Rui Cai, Lynne Ling, Donald Caspary

## Abstract

Aging and challenging signal-in-noise conditions are known to engage use of cortical resources to help maintain speech understanding. Extensive corticothalamic projections are thought to provide attentional, mnemonic and cognitive-related inputs in support of sensory inferior colliculus (IC) inputs to the medial geniculate body (MGB). Here we show that a decrease in modulation depth, a temporally less distinct periodic acoustic signal, leads to a jittered ascending temporal code, changing MGB unit responses from adapting responses to responses showing *repetition-enhancement*, posited to aid identification of important communication and environmental sounds. Young-adult male Fischer Brown Norway rats, injected with the inhibitory opsin archaerhodopsin T (ArchT) into the primary auditory cortex (A1), were subsequently studied using optetrodes to record single-units in MGB. Decreasing the modulation depth of acoustic stimuli significantly increased repetition-enhancement. Repetition-enhancement was blocked by optical inactivation of corticothalamic terminals in MGB. These data support a role for corticothalamic projections in repetition-enhancement, implying that predictive anticipation could be used to improve neural representation of weakly modulated sounds.

**Key points:**

- Aging has been shown to increase temporal jitter in the ascending acoustic code prompting use of cognitive/attentional mechanisms to help better understand communication-like signals.
- Auditory thalamus receives extensive projections from cortex that are implicated in delivering higher-order cortical computations to enhance thalamic responses.
- The present study modeled aging in young rats by using temporally less distinct stimuli shown to alter the pattern of MGB unit responses from response adaptation to repetition-enhancement. Enhanced responses to repeating less temporally distinct modulated stimuli were reversed when inputs from cortex to auditory thalamus were blocked. Collectively, these data argue that low salience temporal signals engage cortical processes to enhance coding of weakly modulated signals in auditory thalamus.

## Introduction

Speech intelligibility can be maintained in noisy backgrounds and in the aged auditory system by increased use of linguistic/contextual redundancies engaged to substitute for sensory deficits (Warren, 1970; Wingfield, 1975; Peelle & Wingfield, 2016; Pichora-Fuller *et al*., 2016; Anderson *et al*., 2020). For young-adults in cluttered acoustic environments and older individuals affected by age-related hearing loss (presbycusis), higher-order/cortical resources are brought into play to help disambiguate acoustic signals (Shinn-Cunningham & Wang, 2008; Davis *et al*., 2011; Obleser, 2014; Başkent *et al*., 2016; Vaden *et al*., 2016; Pichora-Fuller *et al*., 2017). Peripheral deficits only partially account for the age-related loss of speech understanding (Humes *et al*., 2012; Roque *et al*., 2019). Sensory declines in aging may be simulated in young participants by decreasing the temporal distinctiveness of presented acoustic stimuli either by adding noise or decreasing modulation depth, resulting in a temporally jittered ascending acoustic code showing decreases in envelope-locked responses (Dubno *et al*., 1984; Fitzgibbons & Gordon-Salant, 1994; Pichora-Fuller *et al*., 2007; Dimitrijevic *et al*., 2016; Mamo *et al*., 2016). Studies in non-human primates and rabbits using amplitude modulated stimuli have reported an increased neural jitter by decreasing the modulation depth of amplitude-modulated stimuli (Nelson & Carney, 2007; Malone *et al*., 2010). Recent studies support use of increased top-down predictive resources to help decode challenging sensory stimuli such as in speech-in-noise or less temporally distinct speech (Pichora-Fuller *et al*., 2017; Anderson & Karawani, 2020).

Sensory adaptation has been observed in thalamus and cortex, for all sensory modalities, with declining responses for repeated stimuli (Ulanovsky *et al*., 2003; Bartlett & Wang, 2005; Pérez-González & Malmierca, 2014). In contrast to sensory adaptation, repetition-enhancement, perhaps prediction, to a repeating stimulus has been reported when acoustic signals were less temporally distinct, attended to, expected for statistical regularities, and/or with stimuli presented at higher rates in challenging conditions (Luce & Pisoni, 1998; Heinemann *et al*., 2011; de Gardelle *et al*., 2013; Müller *et al*., 2013; Kommajosyula *et al*., 2019). The current study was designed to examine the role of corticothalamic/top-down projections to medial geniculate body (MGB) in mediating repetition adaptation/enhancement responses to repeating stimuli of different modulation depths.

The auditory thalamus is a key subcortical structure suggested to play a critical role in auditory processing. Sensory systems show attention/task/context-dependent changes in thalamic activity, likely reflecting increasingly engaged corticofugal circuits (von Kriegstein *et al*., 2008; Saalmann & Kastner, 2011; Diaz *et al*., 2012; Mihai *et al*., 2019; Tabas & von Kriegstein, 2021). The MGB receives top-down/corticofugal information from extensive descending corticothalamic (CT) projections (Rouiller & Welker, 1991; Winer *et al*., 2001; He, 2003; Bartlett, 2013; Guo *et al*., 2017; Parras *et al*., 2017). These excitatory CT projections originate from cortical layer 5&6 neurons and terminate on the distal dendrites of MGB neurons in all subdivisions, including the lemniscal ventral division and the non-lemniscal dorsal and medial divisions (Bartlett *et al*., 2000; Winer *et al*., 2005; Smith *et al*., 2007). Additionally, MGB receives state and salience-related information from serotonergic/noradrenergic and cholinergic projections (McCormick & Pape, 1990; Sottile *et al*., 2017; Schofield & Hurley, 2018). MGB neurons show stimulus specific adaptation (SSA) to repeated identical stimuli, which upon presentation of an oddball signal show a significant mismatch signal, thought to code for deviance detection and prediction error (Anderson & Malmierca, 2013; Malmierca *et al*., 2015; Parras *et al*., 2017). MGB unit responses show altered tuning and gain changes with manipulation of the auditory cortex/corticofugal influences (Orman & Humphrey, 1981; He, 2003; Tang *et al*., 2012; Malmierca *et al*., 2015). A recent study by Guo *et al*. (2017) showed increased detection of acoustic signals involving CT projections, and CT projections have been shown to be involved in the processing of complex auditory stimuli (Ono *et al*., 2006; Rybalko *et al*., 2006; Homma *et al*., 2017). However, little is known about how CT inputs can alter MGB response properties to repeating signals. The aim of the current study is to examine the impact corticothalamic inputs have on the coding of random vs. repeating sinusoidal amplitude-modulated (SAM) stimuli of differing modulation depths.

Previous MGB single unit studies found that age- and decreased temporal precision (decreased modulation depth or adding noise to the envelope) of the temporal cue significantly increased MGB unit preference (discharge-rate) for repeating SAM stimuli (Cai *et al*., 2016b; Kommajosyula *et al*., 2019). Repetition-enhancement was absent in single-units recorded from MGB in anesthetized rats, suggesting that anesthesia affected thalamic and cortical responses to abolish repetition enhancement (Cai *et al*., 2016b). Collectively, these findings suggest that temporally less distinct acoustic cues and variability due to aging engage top-down/corticofugal influences to enhance responses evoked by a repeating, weakened ascending temporal code. The present study examined MGB single unit responses to determine if increased preference for a repeating less temporally distinct SAM stimulus could be reversed by CT blockade in young, awake rats.

## Materials and Methods

Male Fischer 344 x Brown Norway (FBN) rats (n = 7), aged 4-6 months old, obtained from the NIA Aging Rodent Resource Colony supplied by Charles River, were individually housed on a reverse 12:12-h light-dark cycle with *ad libitum* access to food and water. FBN rats have a long life-span and lower tumor load than other commonly used rat aging models. They have been characterized as a rat model of aging (Cai *et al*., 2018), and age-related changes in central auditory structures have been extensively studied (Caspary *et al*., 2008; Caspary & Llano, 2018; Mafi *et al*., 2020). Procedures were performed in accordance with guidelines and protocols approved (Ref. No. 41-018-004) by the Southern Illinois University School of Medicine Lab Animal Care and Use Committee.

### Microinjection

Adenoviral vectors (AAV-CAG-ArchT-GFP, AAV serotype 1) with light-activated proton pump and eYFP expressed under the control of a CAG (CMV enhancer, chicken beta-Actin promoter and rabbit beta-Globin splice acceptor site) were obtained from the University of North Carolina Vector Core (Chapel Hill, NC). Young-adult FBN rats were anesthetized initially with ketamine (105 mg/kg)/xylazine (7 mg/kg) and maintained with isoflurane (0.5–1%) throughout the duration of the surgery. A small hole was drilled into the skull and dura mater removed. Viral vectors were injected intracranially into left auditory cortex using the Neurostar stereotaxic drill and injection system (stereodrive 015.838, injectomate IM28350, stereodrill DR352; Neurostar, Germany). Coordinates of the injection sites were primary auditory cortex (A1) layers 5 and 6 (L5 and L6), entry at 22° angle laterally (−8.93, −1.8, 4.37 mm relative to bregma). Animals were allowed to recover for 21 days to allow viral expression to transport to the level of CT terminals in the MGB (Fig. 1A).

**Fig. 1.**
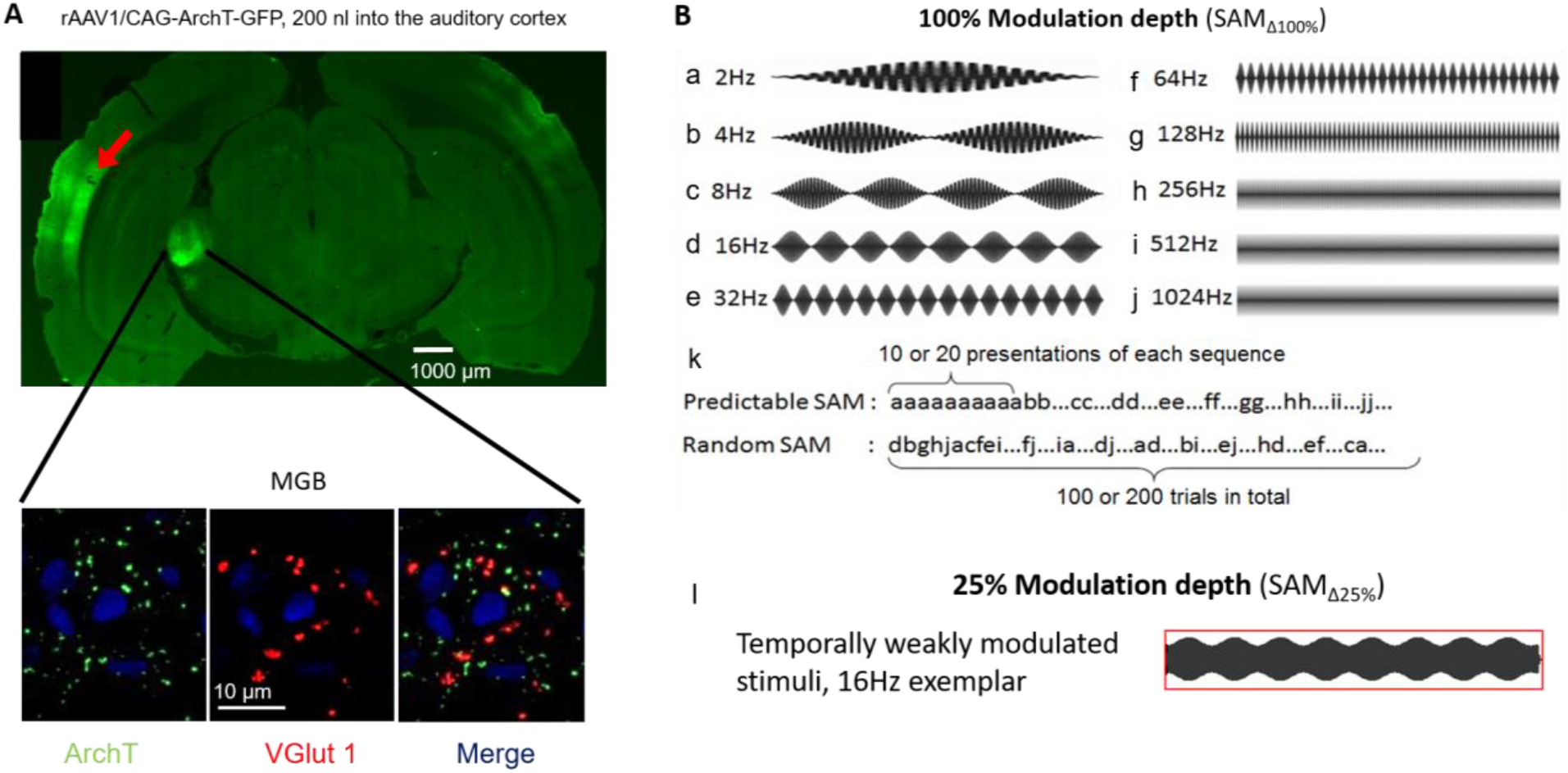
Targeting corticothalamic projections and acoustic stimuli. **A:** Confocal image showing a wide-field and inset of Al GFP-labeled (green) viral injection site and excitatory corticothalamic (CT) projection expressing the ArchT pump. Insets show MGB neurons (63x) receiving labeled projection terminals (ArchT, green), and labeled with glutamatergic marker (VGIutl, red) as well as the nuclear marker (DAPI, blue). Merged image depicts colocalization of ArchT with VGIutl. **B:** Sets of sinusoidally amplitude modulated (SAM) stimuli used in the present study. Standard (100% modulation depth [SAM_Δ100%_]) SAM stimuli with either a tone or broadband noise carrier in 500 ms epochs from 2 Hz to 1024 Hz modulation frequencies [f_mods_] (**B, a-j**)-Stimuli were presented at between 2 Hz to 1024 Hz as either predictable/repeating or random sets (**B, k**). Exemplar waveforms of temporally weakly modulated/less distinct SAM (25% modulation depth [SAM_Δ25%_]) at 16 Hz (**B, I**).

### Acoustic brainstem response (ABR) recording

To ensure normal hearing thresholds, prior to optetrode implantation and 14-21 days after microinjection, auditory brainstem responses (ABR) were collected from all rats as previously described (Wang *et al*., 2009; Cai *et al*., 2016b).

### Awake recordings

Three days following ABR testing, rats began 6-10 day acclimation training in a modified Experimental Conditioning Unit (ECU; Braintree Scientific, Braintree, MA) with free access to water and food reward (1/4 to 1/2 Froot™ Loop) until they could remain quiet/still for up to 3 hours. Prior to surgical implantation, VersaDrive8 optical tetrode drives (Neuralynx, Bozeman, MT) with an additional drive shaft for optical probe were assembled and loaded similarly to VersaDrive4 previously described (Richardson *et al*., 2013; Kalappa *et al*., 2014; Cai *et al*., 2016b). In a dark sound proof booth, there were no other known distractors to divide the rat’s attention during this passive listening task, with SAM stimuli presented from a speaker located above the rat’s head. We recorded 20-25, 45 minute-sessions from each rat. After isolation of a single-unit, spontaneous activity, rate-level functions, and response maps were collected before collecting unit responses to SAM stimulus set. Of the 80 units studied, 95% were clearly isolated single-units (high signal-in-noise ratio, similar amplitude and shape as single units or sorted using principal component analysis) the remaining 5% of units were from small inseparable unit clusters (2-3) are included since no differences in response properties were observed.

All recordings were completed within a 4 week period following implantation recovery. When recordings were complete, rats were anesthetized with ketamine and xylazine as described above and current pulses (5-10 μA for 5 s, nano Z, Neuralynx, Bozeman, MT) were passed through the tips of each tetrode wire, producing a small electrolytic lesions. Rats were cardiac perfused with phosphate-buffered saline (0.1 M, pH 7.4) followed by 4% paraformaldehyde (Sigma, St. Louis, MO), brains were removed, post-fixed for 24 h in 4% paraformaldehyde at 4°C, transferred to 20% sucrose and stored at 4°C until sectioned. To assess the position of recordings, frozen coronal sections (30–35 μm thick) were slide mounted with electrode tracks and lesion sites visible using phase-contrast microscopy. Based on each recording site relative to the final location of the tetrode tip, dimensions of the optetrode placement and MGB anatomy, an approximate location of each recorded unit was derived (Paxinos & Watson, 1998).

### Electrophysiological recordings and optical stimulation

Stimulus paradigms and single unit sorting/recording procedures were the same as for awake rats as in previous studies (Kommajosyula *et al*., 2019). Briefly, extracellularly recoded single spikes, signal to noise ratio of at least 10:1, and with similar waveform were isolated/threholded with small spike unit clusters sorted using of principal component analysis. Stimulus presentation real-time data display and analysis used ANECS software (Dr. K. Hancock, Blue Hills Scientific, Boston, MA). Acoustic signals were generated using a 16-bit D/A converter (TDT RX6, TDT System III, Tucker Davis Technologies, Alachua, FL), and transduced by a Fostex tweeter (model FT17H, Fostex, Middleton, WI) placed 30 cm above animal’s head. The Fostex tweeter was calibrated off-line using a ¼ inch microphone (model: 4938; Brüel & Kjær, Naerum, Denmark) placed at the approximate location of the rat’s head. ANECS generated calibration tables in dB sound pressure level (SPL) were used to set programmable attenuators (TDT PA5) to achieve pure-tone levels accurate to within 2 dB SPL for frequencies up to 45 kHz. The TDT generated “sync-pulse” was connected to an LED optical system (200 µm, 0.39 NA, Thorlabs Inc., NJ) with LED driver (M565F3, LEDD1B, Thorlabs Inc.). Optical stimuli from LED driver were calibrated prior to experiments using optical power meter (S121C and PM121D, Thorlabs Inc., NJ). Optical stimuli were 565 nm wavelength as determined to be the best wavelength for photo-inhibition mediated by ArchT (Han *et al*., 2011). Optogenetic stimulus parameters were chosen to allow for simultaneous stimulation of sound and optical stimuli based on previous and our own preliminary studies: 2.56 mw (∼20.38 mW/mm^2^) intensity presented for 20-40 ms and at 10 Hz regardless of modulation frequencies (*f*_mod_) (Kato *et al*., 2017; Natan *et al*., 2017; Bigelow *et al*., 2019).

### Experimental design: SAM stimulus paradigms and data acquisition

The present study compared the single unit responses in response to three paradigms presented in either a random or repeating paradigm:1) Fully modulated SAM (SAM_Δ100%_), considered the standard clear temporal signal; 2) SAM at 25% modulation depth (SAM_Δ25%_) considered a less temporally distinct signal; 3) SAM_Δ25%_ with during corticothalamic blockade (+ CT blockade) (Fig.1B & 2). There were only small differences (< 2 dB) in total energy levels between the standard (SAM_Δ100%_) and lower modulation depth SAM_Δ25%_ stimuli. We will interchangeably use standard (SAM_Δ100%_) and less temporally distinct SAM (SAM_Δ25%_) across the manuscript. The less temporally distinct SAM stimulus was chosen, in part, as a surrogate for aging to reproduce prior results (Cai *et al*., 2016a; Kommajosyula *et al*., 2019). Kommajosyula et al. (2019) found that SAM_Δ100%_ with1.0kHz noise jittering the envelope gave similar results to SAM_Δ25%_. The SAM carrier was generally BBN, but the unit’s (characteristic frequency) CF was used as carrier if the unit was more strongly driven by CF-tones. Rate modulation transfer functions (rMTFs) and temporal modulation transfer functions (tMTFs) were collected at 30-35 dB above CF or BBN threshold. SAM stimuli were of 450 ms duration, presented at 2/sec with a 4 ms raise-fall; *f*_mods_were stepped between 2 and 1024 Hz (Fig. 1B). SAM stimuli were presented as two separate sets: pseudorandomly, from now on referred to as random across trial (interleaved) *f*_mods_ or identical repeating/blocks of SAM, with each *f*_mod_ repeated (10 times) before being stepped to the next *f*_mod_ in a stepped increasing order (Fig. 1B). To control for order of presentation during repeating trials, we tested *f*_mods_ stepped in descending steps/reverse order, from 1024 to 2 Hz and found that presentation order (descending or ascending) made no difference on spike count. All reported data for repeating SAM trials were stepped from 2 to 1024 Hz. Spikes were collected over a 500 ms period following stimulus onset, with 10 stimulus repetitions at each envelope frequency. Responses to CT blockade examined the role of CT MGB projection during SAM_Δ25%_ stimuli. The effect of CT blockade on coding SAM_Δ100%_ was collected from a subset of MGB units neurons. Data were collected every day for 3-4 weeks after implantation. Data were recorded only if single-unit responses were repeatable and consistent across multiple trials.

Rate-level functions and spontaneous activity (250 epochs of 250 ms each) were recorded in presence and absence of optical blockade. Broadband noise (BBN) (200 ms, 4 ms rise-fall, 2/sec) stimuli were stepped in rate-level functions (0 dB to 80 dB) and responses were collected over a 500 ms period. Response maps were used to determine the CF of sorted single units (Cai & Caspary, 2015). Real-time single unit activity was sampled at 100 kHz and archived for off-line analysis.

### Immunohistochemistry

Free-floating slices were processed in parallel and treated with 0.2% Triton-X for 1 h and incubated for 2 h in blocking solution containing PBS with 0.1% Triton-X, 1.5% normal donkey serum and 3% bovine serum albumin. Sections were transferred to primary antibody solution containing monoclonal mouse anti-vesicular glutamate transporter 1 (VGlut1) antibody (1:750; Millipore, Burlington, MA) in blocking buffer and incubated overnight at room temperature. After washing in PBS, sections were incubated with secondary antibody as follows: donkey anti-mouse IgG (Alexa Fluor 647, 1:150, Jackson ImmunoResearch, West Grove, PA) for 1 h at room temperature. As a negative control, the primary antibody was omitted. Sections were mounted onto slides, cover slipped with VectaShield (Vector Laboratories) and imaged with a Zeiss LSM 800 confocal microscope. Injection of Arch T virus into deep layers of auditory cortex led to expression of GFP tagged ArchT within 4 weeks in the CT terminals at the level of medial geniculate body, as shown by colocalization (yellow) (Fig. 1A).

### Statistical data analysis

Data were collected for MGB single units with SAM_Δ100%_ or SAM_Δ25%_ and CT-blockade as between subject variables. Normality assumptions were met and ANOVA was run to determine significance at the *p* < 0.05 level. Bonferroni corrections were utilized for pairwise comparisons to maintain a type I error level of 5% or less.

Responses were analyzed offline. Phase locking ability was evaluated by the standard vector strength (VS) equation: 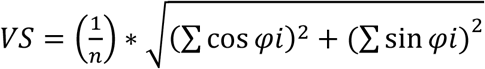 where *n* = total number of spikes and *φi* = the phase of observed spike relative to modulation frequency (Goldberg & Brown, 1969; Yin *et al*., 2011). Statistical significance was assessed using the Rayleigh statistic to account for differences in the number of driven spikes, with Rayleigh statistic values greater than 13.8 considered to be statistically significant (Mardia & Jupp, 2000) (Fig. 2). To compare number of units showing phase locking, a Wilcoxon test was used followed by a Bonferroni correction for multiple comparisons.

**Fig. 2.**
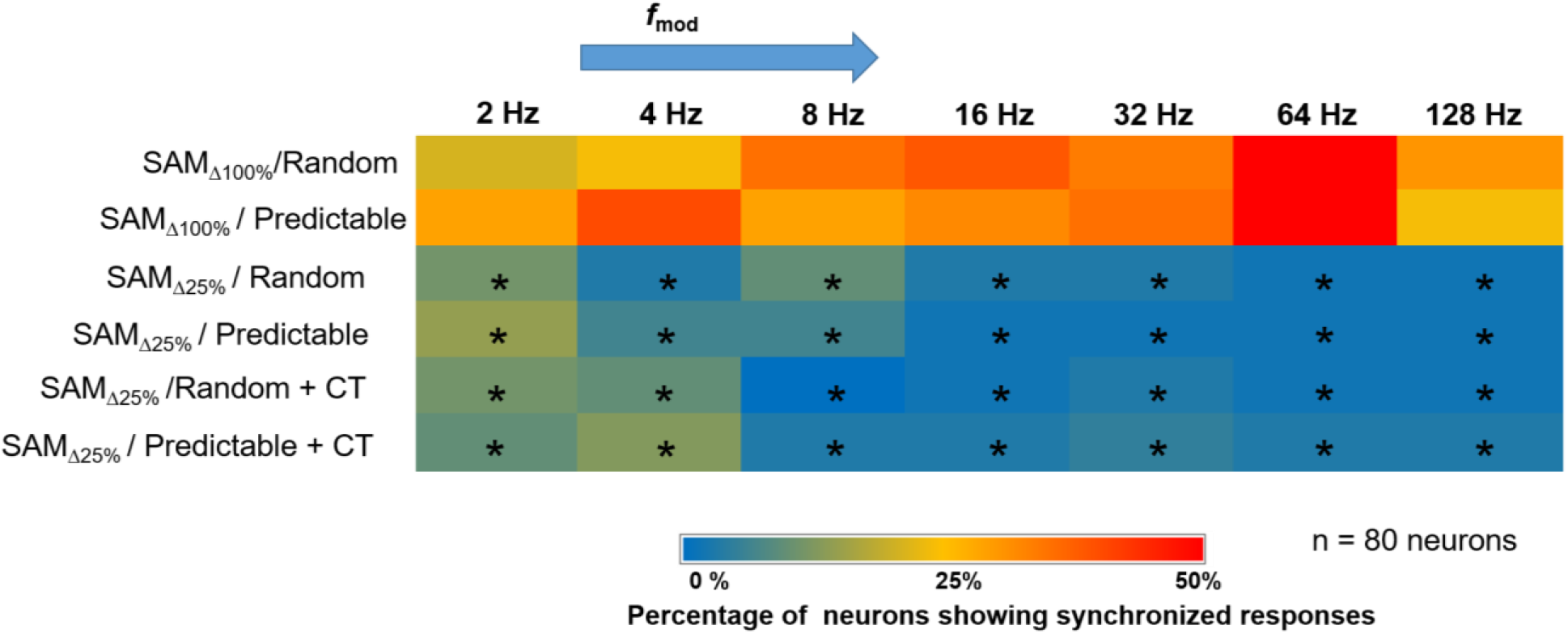
Effects of stimulus modulation depth on temporal locking properties of MGB units: To assess the ability of units to temporally follow the SAM stimulus, the Rayleigh score for each *f*_mods_ (2-128) was used to generate a heat map based on the temporal responses of all 80 MGB units studied. MGB units might lock to a single or multi *f*ms based on the Rayleigh score. Warmth of color indicates the percentage of neurons (out of 80) showing temporal-locking (Rayleigh statistic ≥ 13.8) to the SAM stimuli. Hot colors (red) indicate a higher percentage of units showing temporal-locking (e.g SAM_Δ100%_at 64 Hz *f*m), whereas cool colors (blue) indicate a lower percentage of units showing temporal locking (e.g. SAM_Δ25%_at 16 Hz *f*m). Significant differences were observed between SAMΔ100_%_ and regardless of order of presentation, with and without CT blockade (Wilcoxon test followed by Bonferroni correction, *p* < 0.05).

Rate-level functions determined using spike rate in response to BBN were quantified across intensities and compared between control and CT blockade paradigms using repeated measures ANOVA with Bonferroni correction. Spontaneous activity measured using spike rate across 250 ms epochs in 10 ms bins were compared between control and CT blockade paradigms using repeated measures ANOVA with Bonferroni correction. Preliminary analysis involved differences between order of presentation and across stimulus conditions using total spike counts from 10 trials at 10 different *f*_mods_. Differences between orders of presentation were compared across random or repeating presentation of stimuli between SAM_Δ100%,_ SAM_Δ25%,_and SAM_Δ25%_+CT blockade condition using repeated measures ANOVA followed by post-hoc Bonferroni corrections.

Differences between stimulus conditions were compared using a preference ratio (PR) calculated across all *f*_mods_ (PR = total spikes in repeating trials/total spikes in random trials). A ratio smaller than 0.95 suggests the unit is a random preferring unit; a ratio larger than 1.05 suggest the unit is repetition preferring unit; while a ratio between the range of 0.95 and 1.05 were considered non-selective units (Fig. 3). The rationale for use of 10 % change in firing as a criteria was based on previous studies (Ghitza *et al*., 2006; Cai & Caspary, 2015; Cai *et al*., 2016b). Chi-Square test was used to compare the PR across conditions.

**Fig. 3.**
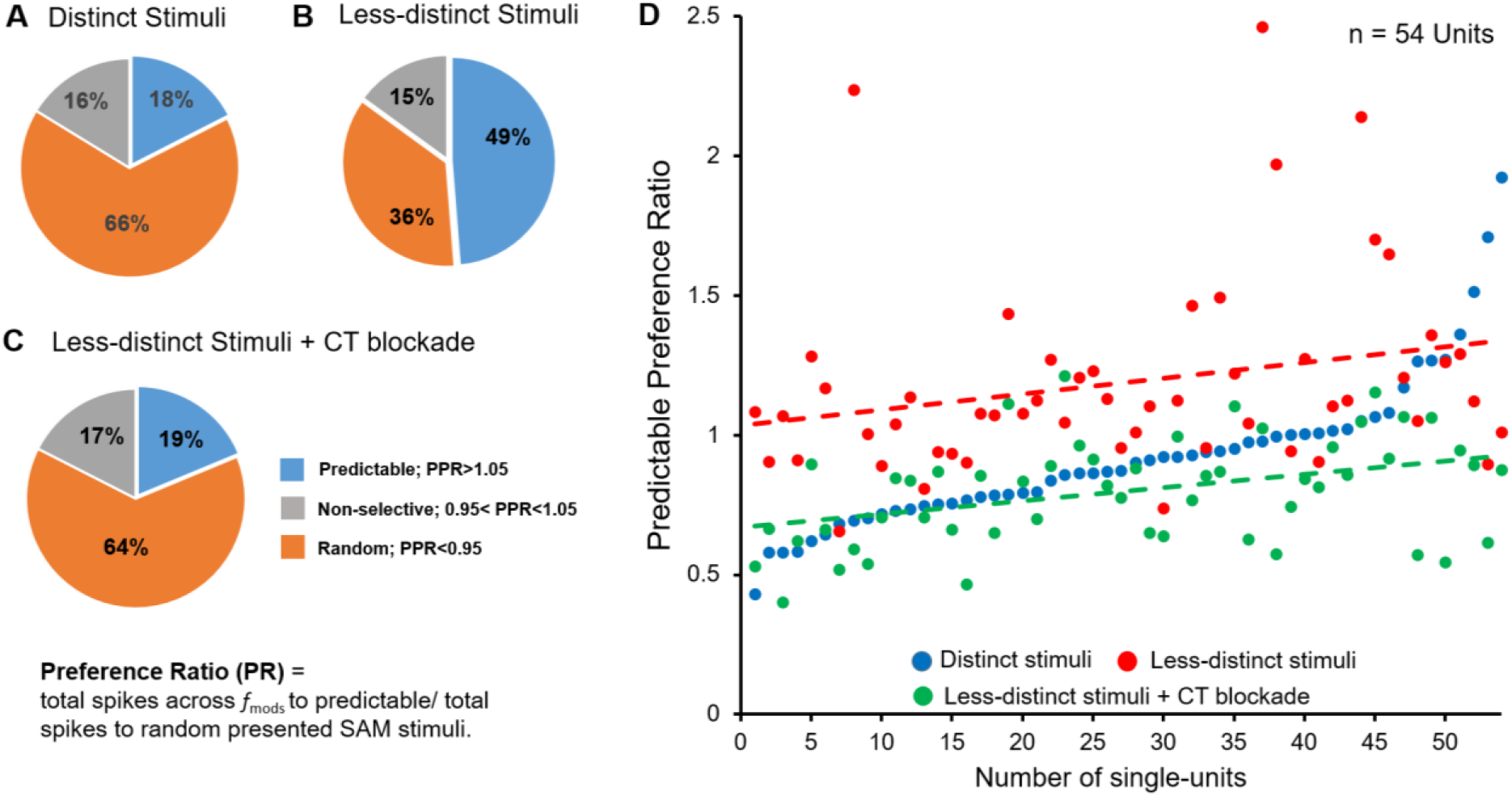
Random vs. predictable/repetition preference with and without CT blockade. Preference ratios (PR) (total spikes to predictable trials/total spikes to random trials) across all *f*_mods_ in response to distinct, less distinct SAM stimuli, less-distinct stimuli with corticothalamic blockade (CT blockade). **A**: Unit recording from awake rat MGB showed a clear preference for random distinct SAM_Δ100%_ stimuli. **B**: Responses to predictable (repeating) SAM stimuli increased from 18% (14/80). to 49% (39/80). in response to SAM_Δ25%_ across *f*_mods_ **C**: Optical CT blockade reversed the predictable preference of MGB neurons to 19% (14/80. in response to less SAM_Δ25%_ SAM Significant differences were seen between SAM_Δ100%_.,vs. SAM_Δ25%_. SAM_Δ25%_ VS. SAM_Δ25%_ + CT blockade and SAM_Δ25%_ + CT blockade vs. SAM_Δ25%_ + recovery (Chi-Square test, p < 0.05). D: PR values plotted on a continuum of increasing PPI values for each of 54 MGB units showing differential responses to distinct, SAMi_Δ100%_ (blue dots) vs. less-distinct, SAM_Δ25%_stimuli (red dots) and SAM_Δ25%_ with CT blockade (green dots). The green trend line shows that CT blockade dramatically decreased the PR in response to SAM_Δ25%_ (red trend-line) approaching the response to SAM_Δ100%_ stimuli (blue dots).

**Fig. 4.**
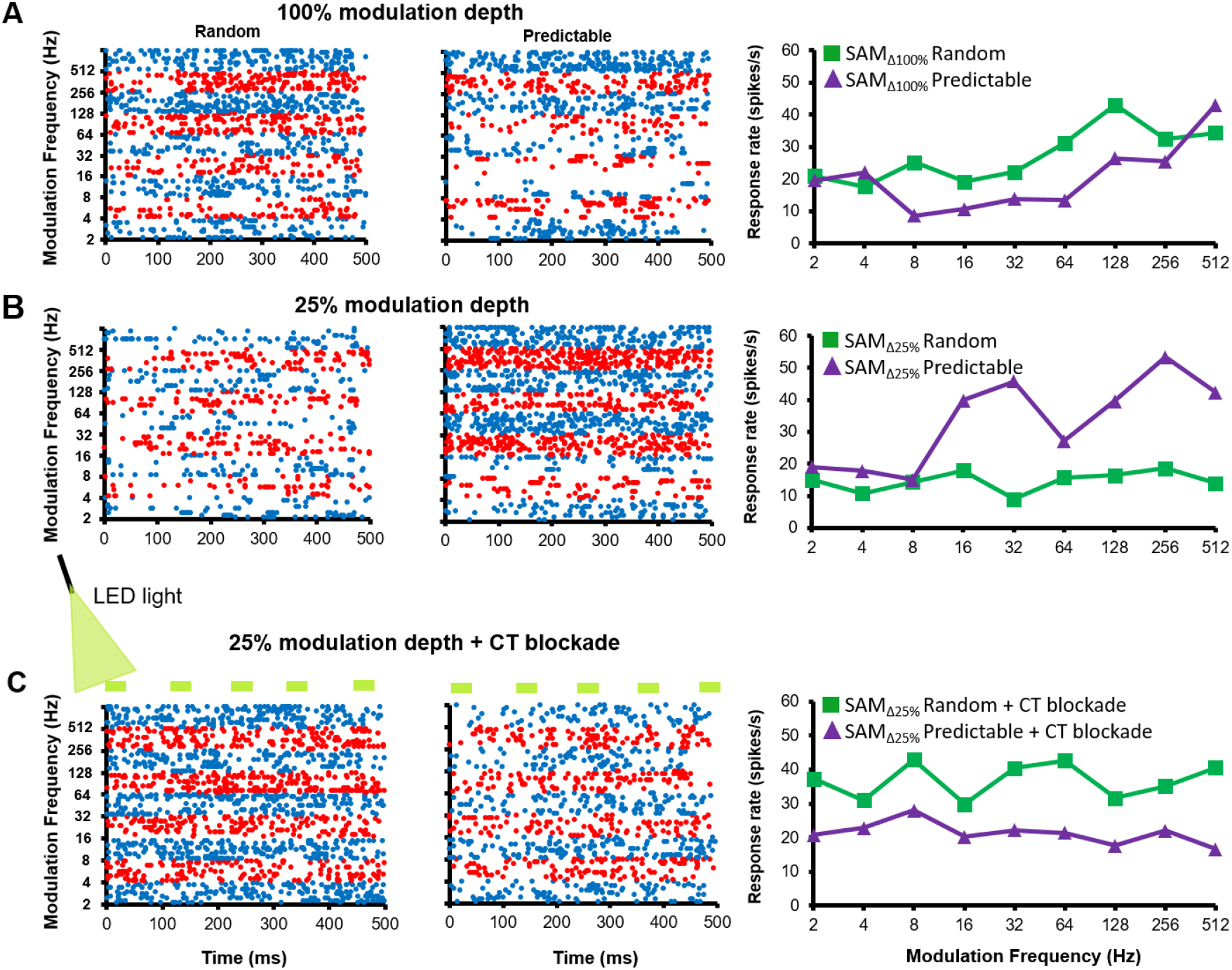
Exemplar MGB unit showing differential responses to SAM presentation order, modulation depth and CT blockade: **A.** a representative MGB unit showing a higher discharge rate (spikes/sec) to randomly presented SAM_Δ100%_ across *f*_mods_ than to predictable/repeating SAM_Δ100%_ stimuli in dot raster and rate-modulation transfer functions (rMTFs). **B**. When modulation depth was decreased to SAM_Δ25%_, less distinct stimuli, the same MGB unit showed mcreased/greater responses to a predictable/repeating SAM, especially at higher fms. **C**. Optical blockade of CT input resulted in a return to strong random preference even in response to less distinct stimuli, SAM_Δ25%_ in this same exemplar.

Modulation transfer functions (MTFs) were determined using spike rate (rMTF) measurements at each *f*_mod_ tested. The rMTF data were used for further quantitative analyses. A predictable preference index (PPI) was calculated using the area under the curve (AUC) and the equation: PPI = [(AUC_REP_-AUC_RAN_)/ (AUC_REP_+AUC_RAN_)], modified from the novelty response index (Lumani & Zhang, 2010; Cai *et al*., 2016b). The area under successive frequency segments of the rMTF curve (AUC) values were based on rMTF curve calculated using GraphPad Prism. The range of PPI values varied between -1 to +1: +1 represented a repetition preferring unit response, and -1 represented a random preferring unit response (Figs. 5 and 6). By calculating the AUC for specific *f*_mod_ ranges, changes between sets of *f*_mod_ could be compared. Repeated-measures ANOVA followed by post-hoc Tukey correction for multiple comparisons was used to compare PPI values.

**Fig. 5.**
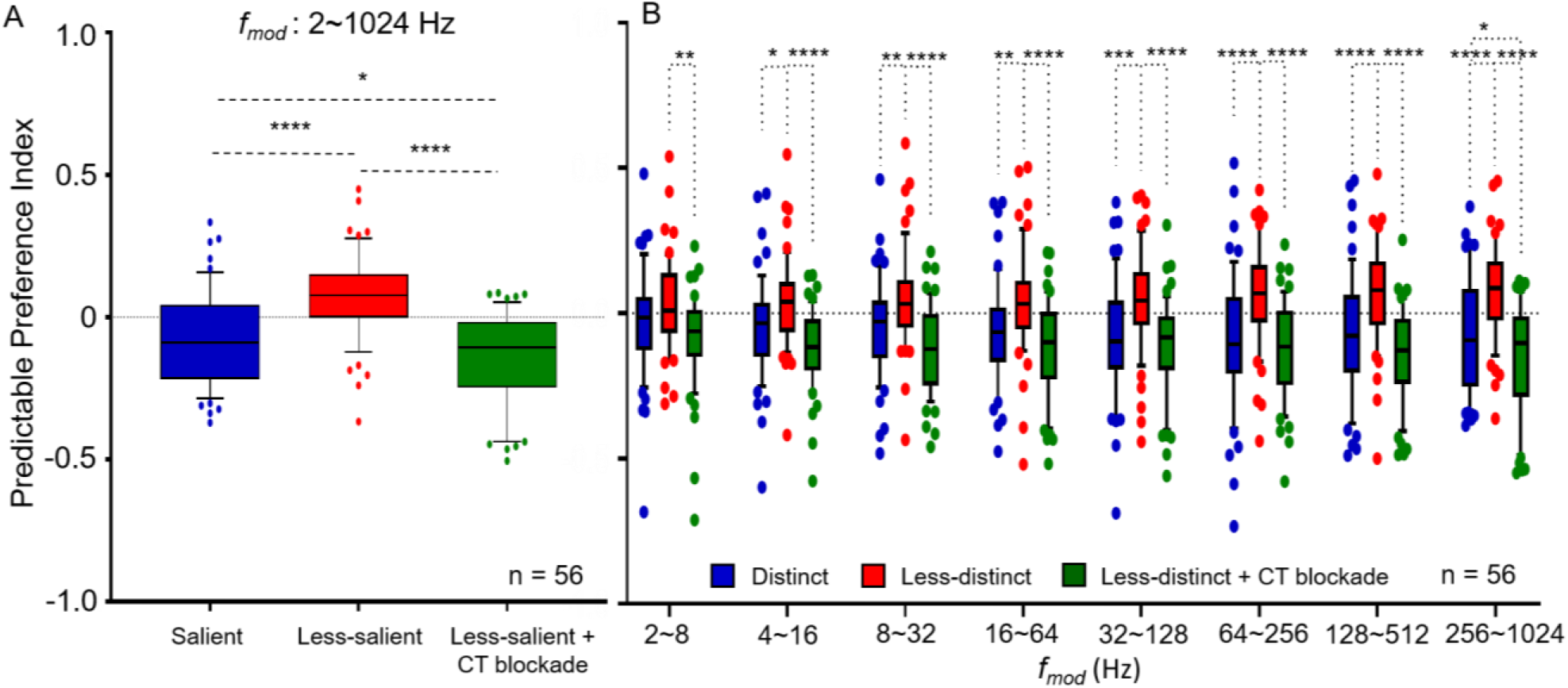
Predictable preference index (PPI) for MGB unit’s sensitive to stimulus depth of modulation: PPI’s were calculated (see text) for MGB responses to random and predictable trials across all *f*_mods_ combined and for specific subsets of *f*_mods_ **A.**. For all *f*_mods_ combined, MGB units (n = 56) showed significant increases in PPI values (red bar) when switching from SAM_Δ100%_ to less distinct SAM_Δ25%_ stimuli (blue bar). The observed increase in PPI was reversed (green bar) with corticothalamic (CT) blockade. **B.** PPI values for MGB neurons showed significantly increased PPIs to SAM_Δ100%_ especially at higher *f*_mods_ with CT blockade reversing these increases. (Data are presented as the mean ± SEM; repeated-measures ANOVA followed by post *hoc* Tukey s correction were used for analyses (Graphpad), \**p* < 0.05; * \**p* < 0.01; * * \**p* < 0.001; * * * \**p* < 0.0001.

**Fig. 6.**
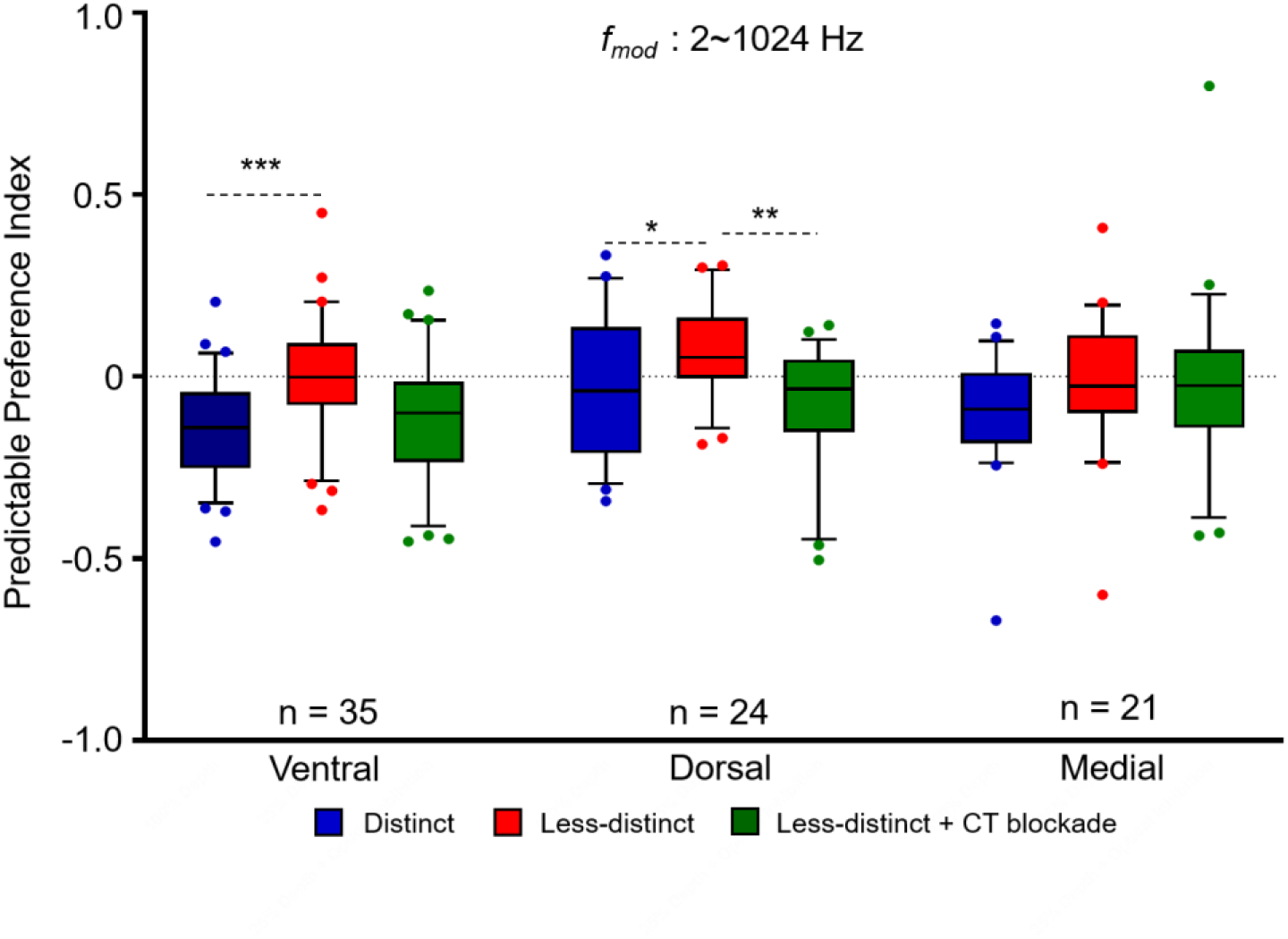
MGB region specific changes in predictable preference index (PPI) for unit’s sensitive to stimulus depth of modulation. PPI’s were calculated (see text) for MGB units located in the three major divisions of the MGB. Responses to random vs. predictable SAM across all *f*_mods_ combined with and without CT blockade. Across *f*_mods_ dorsal (24) and ventral (39), MGB units showed significant increases in PPI values (red bar) when switching from SAM_Δ25%_ to SAM_Δ100%_ Corticothalamic (CT) blockade reversed this significant increase for dorsal and ventral MGB units. These changes were not observed in the medial division. Data are presented as the mean ± SEM; repeated-measures ANOVA followed by *post hoc* Tukey’s correction were used for analyses (Graphpad). \**p* _<_ 0.05; * \**p* _<_ 0.01; * * \**p* < 0.001.

Trial-to-trial responses to repeating/predictable SAM presentation showed repetition-enhancement at temporally challenging (higher frequency) *f*_mods_ (*f*_mods_ 128 Hz*-*1024 Hz) (Cai *et al*., 2016b; Kommajosyula *et al*., 2019). Differences in firing rate trend-line slopes between the three groups (standard SAM were compared using two-tailed ANCOVA, followed by Friedman test with a post-hoc Wilcoxon test to analyze spike rate differences at each trial (Fig. 7).

**Fig. 7.**
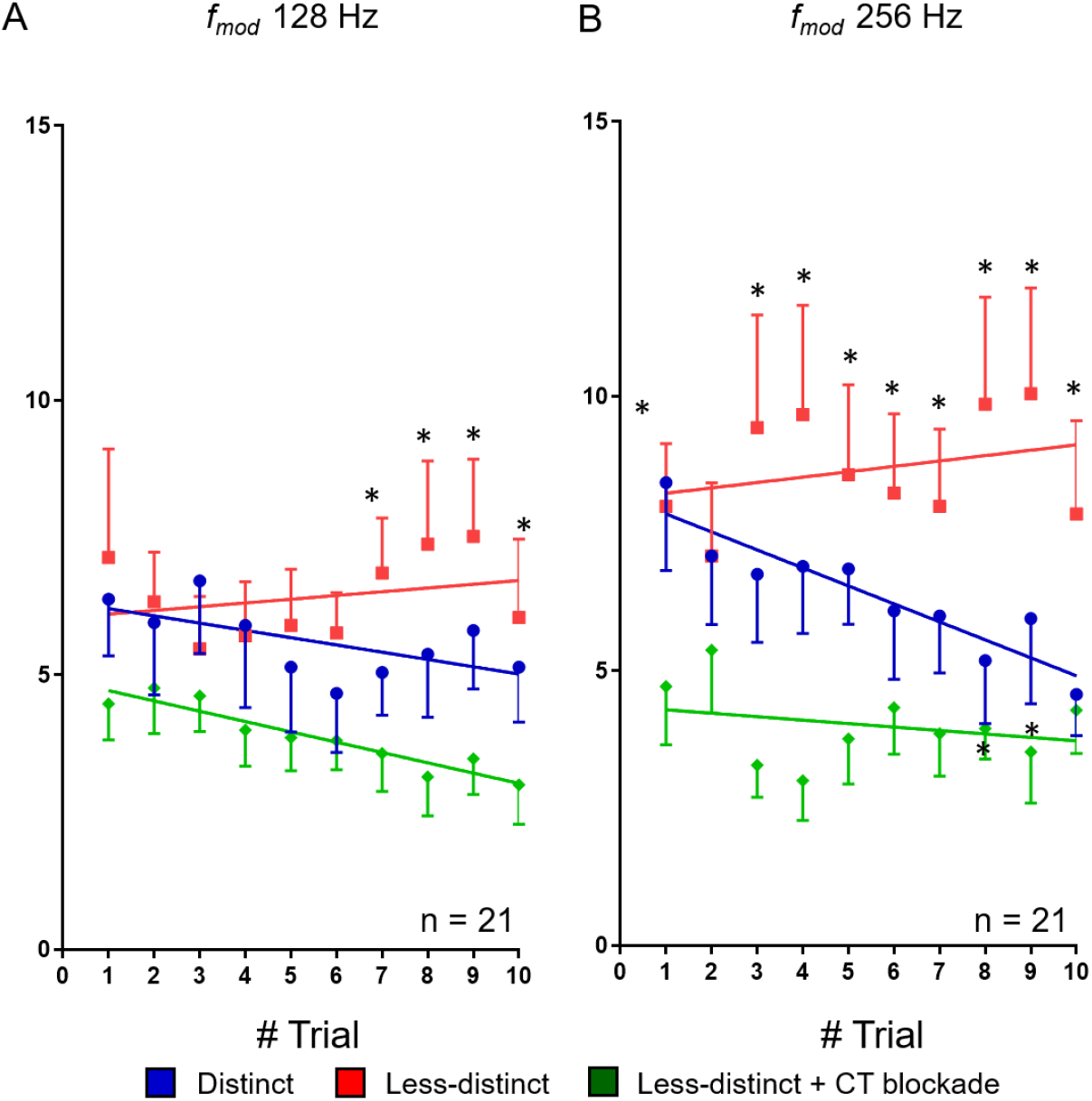
Trial-by-trial response analysis to SAM _Δ100%_ to SAM _Δ25%_ with and without corticothalamic (CT) blockade. Single-units showing PPI changes larger than 0.3 at high f_mods_ when switch from SAM_Δ100%_ to *SAM*_Δ25%_ are included in the trail by trial analysis. Group (n = 21) trial-by-trial responses to predictable SAM at f_mods_ 128Hz (A) and 256Hz (B). These units show adapting responses to 10 presentations of repeating salient SAM_Δ100%_ stimuli (blue dot). Decreasing SAM modulation depth switched the trial-by-trial responses from adapting to predictable with spikes increasing with each successive presentation of the SAM_Δ25%_ stimulus (red dot). Optical CT blockade reversed the predictive response (green dot). Trend line slopes were significantly different for the three conditions for average spikes to predictable presentation of at f_mod_ 128 Hz (A, ANCOVA, two-tailed, p _<_ 0.05). Differences were significant at individual trial 7, 8, 9 and 10 in between SAM_Δ25%_ and *SAM*_Δ25%_ CT stimulus conditions (p _<_ 0.05, Friedman test followed Wilcoxon test) (A). Similarly, Trend line slopes were significantly different for the three conditions for average spikes to predictable presentation at fmod 256 Hz (B) (ANCOVA, two-tailed, p < 0.05). Differences were significantly different at trial 1, 3, 4, 5, 6, 7, 8, 9, and 10 between SAM_Δ25%_ vs. *SAM*_Δ25%_ with CT blockade. There were significant differences between SAM_Δ100%_ and SAM_Δ25%_ stimuli at trial 8 and 9 in their firing rates (B) (p < 0.05, Friedman test followed Wilcoxon test).

Repeated measures ANOVA followed by post-hoc Bonferroni corrections were used to test statistical significance. Statistical analysis was performed using GraphPad Prism 6 and IBM SPSS version 24. All values are expressed as means ± SEM. ^*^*p* < 0.05, ^**^*p* < 0.01, ^***^*p* < 0.001, ^****^*p* < 0.0001, were treated as statistical significance level.

## Results

Eighty MGB units, responding to sinusoidal amplitude modulation stimuli (SAM) were recorded from the MGB in awake, passively listening, young-adult FBN rats. Consistent with previous studies, MGB single-unit responses to SAM stimuli showed band-pass, low-pass, high-pass, mixed or atypical rMTFs, showing synchronized and asynchronized or mixed responses (Bartlett & Wang, 2007).

### Basic response properties with CT blockade

There were no significant changes in spontaneous activity with CT blockade compared to control condition (13.85 ± 1.27 vs 13.26 ± 1.34, n = 45; *p* = 0.282). Rate-level functions showed significant decreases in responses across intensities with CT blockade compared to control (Multivariate ANOVA, *p* = 0.040) with significant differences for comparisons at a couple of intensities (Table 1).

**Table 1:** Rate-level functions under control and CT blockade conditions for 60 units.

| Intensity (dB) | Control<br>(Mean±SEM) | CT blockade<br>(Mean±SEM) | p-value* |
| --- | --- | --- | --- |
| 0 | 15.927±1.5 | 15.493±1.5 | 0.564 |
| 10 | 15.75±1.7 | 14.306±1.5 | 0.152 |
| 20 | 15.195±1.5 | 13.941±1.5 | 0.07 |
| 30 | 14.797±1.5 | 13.997±1.5 | 0.346 |
| 40 | 16.0396±1.7 | 13.472±1.4 | 0.004 |
| 50 | 14.854±1.5 | 13.920±1.5 | 0.17 |
| 60 | 15.429±1.5 | 13.791±1.4 | 0.031 |
| 70 | 15.462±1.4 | 15.008±1.5 | 0.437 |
| 80 | 18.062±1.8 | 18.182±1.7 | 0.908 |

Comparisons are made between control (column 2) vs. CT blockade (column 3) in 60 neurons; \*Column 4 represents the Bonferroni-corrected p-values for comparisons between MGB single-unit responses to control and CT blockade.

### Decrease in modulation depth decreases envelope-locking of MGB neurons

Decreasing modulation depth to SAM_Δ25%_ decreased envelope locking of MGB units studied relative to SAM_Δ100%_ stimuli, as measured using the Rayleigh score across *f*_mods_ (2-128 Hz) (Fig. 2). A higher percentage of MGB units showed temporal locking (Rayleigh statistic ≥13.8) to the standard stimuli (SAM_Δ100%_) than to the SAM_Δ25%_ stimuli across *f*_mods_ tested (Table 2). CT blockade did not alter percentages of envelope-locking responses to less-distinct/SAM_Δ25%_ stimuli across *f*_mods_ tested. These data show decreased temporal locking in response to SAM_Δ25%_ stimuli and that temporal locking was relatively independent of top-down modulation. These results are similar to findings showing decreases in temporal locking when adding noise to the SAM periodic envelope (Kommajosyula *et al*., 2019). Here we focus on rate responses of MGB single-units and the effect of CT projections on MGB single-unit response properties.

**Table 2:** Bonferroni-corrected p-values for percentage of envelope-locking units with changing sound stimuli modulation depth.

| fm range (Hz) | SAM<sub>Δ100</sub> vs. SAM<sub>Δ25%</sub>* |  | SAM<sub>Δ100</sub> vs. SAM<sub>Δ25%</sub>*<br>+CT blockade* |  |
| --- | --- | --- | --- | --- |
|  | Random | Repeating | Random | Repeating |
| 2 | 0.037 | 0.0079 | 0.037 | 0.0014 |
| 4 | 0.00082 | 0.000012 | 0.0071 | 0.000098 |
| 8 | 0.00016 | 0.00048 | 0.00000072 | 0.00016 |
| 16 | 0.0000072 | 0.000034 | 0.0000042 | 0.000058 |
| 32 | 0.000034 | 0.000012 | 0.000034 | 0.000034 |
| 64 | 0.0000001 | 0.0000001 | 0.0000001 | 0.00000019 |
| 128 | 0.000058 | 0.00048 | 0.000058 | 0.00082 |

\*Each row represents the Bonferroni-corrected p-values following the Wilcoxon tests for corresponding fm (in Hz) in the first column of the row. Comparisons are made between salient vs. less-salient (column 2) and salient vs. less-salient+ CT blockade (column 3) in all the of neurons (n=80).

### Decreased modulation depth and CT blockade significantly alter MGB unit rate response to random vs. repeating SAM

Total spike counts in response to SAM stimuli presented in random or repeating trials were compared across stimulus sets with and without CT blockade (standard SAM at 100% depth of modulation [SAM_Δ100%_]), less distinct (SAM at 25% depth of modulation [SAM_Δ25%_]), less distinct SAM_Δ25%_ + CT blockade) (Fig. 1B and methods for details). Consistent with Kommajosyula et al. (2019), 66% (56 of 80) MGB units preferred randomly presented SAM_Δ100%_ stimuli (Fig. 3A). When modulation depth was reduced to SAM_Δ25%,_ there was a significant increase in the percentage of MGB units showing a rate preference for repeating stimuli (18% vs. 49%, *X*^2^(4, N = 80) = 88.789, *p* = 2.3812E-18) (Fig. 3A&B). This switch in preference toward repeating less distinct SAM_Δ25%_ was reversed by CT blockade in MGB (49% vs. 19%, *X*^2^(4, N = 80) = 84.884, *p* = 1.6054E-17) (Fig. 3B&C). Following termination of CT optical blockade, MGB unit responses returned to showing increased response preference for repeating less distinct/SAM_Δ25%_ (19% vs. 39%, *X*^2^(6,N = 80) =106.386, *p* = 1.1628E-20, data not shown).

Ninety percent (72/80) of MGB units changed their PRs toward repeated stimuli in response to the switch in modulation depth/CT blockade (change in PR > 0.1). Seventy-five percent (54/72) of those units shifted their preference from repeated back to random stimuli with CT blockade at SAM_Δ25%_. The PR scores for each of the 54 MGB units were plotted on a continuum of increasing PR score for SAM_Δ100%_, with PR for SAM_Δ25%_ (with or without CT blockade) also plotted for each unit (Fig. 3D). PR trend lines show an increase in PR to repeating stimuli when switching from SAM_Δ100%_ to SAM_Δ25%_ for most units (Fig. 3D-red line). CT blockade during SAM_Δ25%_ stimuli (green trend line) returns the PR or preference for random stimuli, to levels which approximate but are below responses for SAM_Δ100%_. Reducing SAM modulation depth increased repetition-enhancement in 54/72 neurons, while CT blockade reversed the switch from repetition-enhancement to adapting responses (Fig. 3D).

The 18 remaining MGB units of the 72 units did not show a change in PR with a decrease in SAM temporal distinctiveness (SAM_Δ100%_ to SAM_Δ25%_) but showed increase in PR, or a preference for repeated stimuli when switched to SAM_Δ25%_ with optical CT blockade. Eight MGB neurons unresponsive to optical blockade were not included in the analysis.

Changes in response to modulation depth and CT blockade are shown for an exemplar MGB unit (Fig.4). Switching to less-distinct SAM_Δ25%_ showed a two-fold increase in responses to repeating trials across a range of modulation frequencies, which was reversed by CT blockade (Fig. 4B&C).

Since PR does not differentiate differences across *f*_mods_, we calculated the predictable preference index (PPI), a quantitative measure derived from area under the curve (AUC) values across groups of modulation frequencies, PPI = [(AUC_REP_-AUC_RAN_)/ (AUC_REP_+AUC_RAN_)]. Higher PPI values indicate increased preference for repeating trials, while lower PPI values indicate a preference for randomly presented trials. PPI values were lower for standard stimuli (SAM_Δ100%_) across all *f*_mods_ tested (Fig. 5A). Seventy-nine percent of MGB units (56/71) showed increased PPI value with decreased modulation depth (SAM_Δ25%_), indicating repetition-enhancement. CT blockade during presentation of SAM_Δ25%_ reversed the notable increase in PPI (repeated measures ANOVA, F(2, 165) = 39.512, *p* = 2.682E-11, Bonferroni corrected p-values (standard vs. less-salient = 0.000001; SAM_Δ25%_ vs. SAM_Δ25%_+ CT blockade = 1.4624E-11; SAM_Δ100%_ vs. SAM_Δ25%_+ CT blockade = 0.019) (Fig. 5A). Changes in PPI were determined for sets of increasing *f*_mods_ across different stimulus groups (Fig. 5B). SAM_Δ25%_ significantly increased PPI values and these changes were more pronounced at higher *f*_mods_. CT blockade significantly decreased PPI values across *f*_mods_ (Fig. 5B). At *f*_mods_ between 256-1024 Hz, PPI values were significantly decreased by CT blockade even when compared to standard, SAM_Δ100%_ stimuli (Table 3 for repeated measures ANOVA, Bonferroni corrected p-values and comparisons at each *f*m range) (Fig. 5B).These results suggest that MGB responses to standard, SAM_Δ100%_ stimuli show a degree of CT influences at the higher *f*_mods_ tested. For 13 single-units, the effects of CT blockade at SAM_Δ100%_was tested in resopnses to sequencial/repeating trails with and without CT blockade. There were no significant differences in spike rates (SAM_Δ100%_ vs. SAM_Δ100%_+ CT blockade = 17.62615 ± 3.52428 vs. 15.2132 ± 2.9107, *p* = 0.0529, T-test) and for PPI values between the two conditions across all *f*_mods_ (SAM_Δ100%_ vs. SAM_Δ100%_+ CT blockade = -0.03926 ± 0.0393 vs. -0.03136 ± 0.0316, *p* = 0.8611, T-test). This results supports the hypothesis that additional top-down resources were engaged by temporally less distinct SAM stimuli.

**Table 3:** Bonferroni-corrected p-values for PPI values of 56 units sensitive to modulation depth change.

| *f*m range (Hz) | SAM<sub>Δ100</sub> vs. SAM<sub>Δ25%</sub>* | SAM<sub>Δ25</sub> vs. SAM<sub>Δ25%</sub>*<br>+CT blockade* | SAM<sub>Δ100</sub> vs. SAM<sub>Δ25%</sub>*<br>+CT blockade* |
| --- | --- | --- | --- |
| 2~1024 | 0.000001 | 1.4624E-11 | 0.019 |
| 2~8 | 0.194306274 | 0.002414952 | 0.238624372 |
| 4~16 | 0.026271845 | 7.2624E-06 | 0.081380638 |
| 8~32 | 0.004712185 | 3.10386E-07 | 0.071969343 |
| 16~64 | 0.004245266 | 3.88949E-06 | 0.213901181 |
| 32~128 | 0.000870169 | 8.76613E-06 | 0.533565168 |
| 64~256 | 5.21613E-05 | 4.47651E-08 | 0.347311451 |
| 128~512 | 6.14401E-05 | 7.89532E-10 | 0.091124673 |
| 256~1024 | 3.8719E-05 | 4.2874E-11 | 0.039640353 |

\*Each row represents the Bonferroni-corrected p-values to corresponding *f*m range (in Hz) in the first column of the row. Comparisons are made between salient vs. less-salient (column 2); less-salient vs. less-salient + CT blockade (column 3); and salient vs. less-salient+ CT blockade (column 4) in the majority of neurons (n=56).

The 15 MGB units that did not show PPI changes in modulation depth paradoxically showed significantly increased PPI values with CT blockade, across *f*_mods_ examined (Table 4).

**Table 4:** Bonferroni-corrected p-values for PPI values of 15 units insensitive to modulation depth change.

| *f*m range (Hz) | SAM<sub>Δ100</sub> vs. SAM<sub>Δ25%</sub>* | SAM<sub>Δ25</sub> vs. SAM<sub>Δ25%</sub>*<br>+CT blockade* | SAM<sub>Δ100</sub> vs. SAM<sub>Δ25%</sub>*<br>+CT blockade* |
| --- | --- | --- | --- |
| 2~1024 | 0.823374867 | 1.48214E-05 | 0.000142022 |
| 2~8 | 0.774652034 | 0.992047279 | 0.840986588 |
| 4~16 | 0.811395141 | 0.79258215 | 0.41545121 |
| 8~32 | 0.973549441 | 0.238930012 | 0.342751939 |
| 16~64 | 0.946456635 | 0.018936675 | 0.044413028 |
| 32~128 | 0.800376342 | 0.002708837 | 0.000256347 |
| 64~256 | 0.239459428 | 0.003792644 | 5.52114E-06 |
| 128~512 | 0.705864677 | 0.001001792 | 4.09608E-05 |
| 256~1024 | 0.997297984 | 0.000232669 | 0.000175392 |

\*Each row represents the Bonferroni-corrected p-values to corresponding *f*m range (in Hz) in the first column of the row. Comparisons are made between salient vs. less-salient (column 2); less-salient vs. less-salient + CT blockade (column 3); and salient vs. less-salient+ CT blockade (column 4) in the minority of neurons (n=15).

### Trial by trial analysis

Based on the PPI results (Fig. 5) suggesting that sensory responses were adapting and top-down MGB inputs caused repetition-enhancement, we examined trial-by-trial data to 10 successive presentations of SAM stimuli, for the 21 MGB units with the highest PPI values (> 0.3) at *f*_mods_ that showed the largest changes (Fig. 7). Group data for repeating presentations of SAM stimuli (128 Hz and 256 Hz *f*_mod_) showed clear adaptation across trials for SAM_Δ100%_, while reducing SAM depth changed the slope to repetition-enhancement. CT blockade reversed the trial-by-trial repetition-enhancement in response to repeating SAM_Δ25%_ stimuli (Fig. 7A&B). Trend line slopes for average spikes were significantly different across the three conditions for repeating presentation at 128 Hz *f*_mod_ (F(2,24) = 4.885. *p =* 0.0166). Differences were significant for individual trials 7, 8, 9 and 10 between less-distinct and less-distinct with CT blockade (Friedman test followed Wilcoxon test and respective *p*–values for each trial are mentioned: (trial 7, *p* = 0.0021; trial 8, *p* = 0.0011; trial 9, *p* = 0.0027; trial 10, *p* = 0.009) (Fig. 7A). Responses to a repeating SAM (*f*_mod_ 256 Hz) significantly adapted to SAM_Δ100%_ stimuli, while increasing responses across trials to SAM_Δ25%_, which was reversed by CT blockade (ANCOVA, two-tailed, *F(2,24) = 6*.*527, p = 0*.*0055*). Differences were significant for all trials but trial 2 between SAM_Δ25%_ to SAM_Δ25%_ with CT blockade (Friedman test followed Wilcoxon test and respective p –values for each trial are mentioned: (trial 1, *p* = 0.006; trial 3, *p* = 0.00018; trial 4, *p* = 0.00046; trial 5, *p* = 0.0002; trial 6, *p* = 0.0018; trial 7, *p* = 0.0034; trial 8, *p* = 0.0013; trial 9, *p* = 0.0004; trial 10, *p* = 0.038)) (Fig. 7B). The same trends were seen for trial-by-trial spike rate comparisons for *f*_mods_ 512 and 1024 Hz. The impact of onset responses on trial-by-trial rate data was examined by removing the first 50 ms. There were no significant differences in these data with or without inclusion of 50 ms onset across the three stimulus conditions (data not shown).

### MGB subdivisions

PPI values across *f*_mods_ were examined for all 80 units based on their location within the major MGB subdivisions (Fig. 6). PPI values were significantly increased in ventral and dorsal MGB when modulation depth was reduced from SAM_Δ100%_ to SAM_Δ25%_ (Fig.6). Corticothalamic blockade reversed the PPI changes in the dorsal division with a trend toward reversal in the ventral MGB (repeated measures ANOVA F(1.714, 132) = 8.562, *p* = 0.0006, Bonferroni corrected p-values across all *f*ms in ventral division (SAM_Δ100%_ to SAM_Δ25%_ = 0.0002; SAM_Δ25%_ to SAM_Δ25%_ + CT blockade = 0.0859; SAM_Δ100%_ to SAM_Δ25%_ + CT blockade = 0.5902); Bonferroni corrected p-values across all *f*ms in dorsal division (SAM_Δ100%_ to SAM_Δ25%_ = 0.0389; SAM_Δ25%_ to SAM_Δ25%_ + CT blockade = 0.0012; SAM_Δ100%_ to SAM_Δ25%_ + CT blockade = 0.5146); Fig. 6). None of these changes were significant in the medial division of the MGB (Bonferroni corrected p-values across all *f*ms in medial division (SAM_Δ100%_ to SAM_Δ25%_ = 0.1541; SAM_Δ25%_ to SAM_Δ25%_ +CT blockade = 0.9971; SAM_Δ100%_ to SAM_Δ25%_ + CT blockade = 0.3117; Fig. 6).

### Spike-rate changes with altered SAM modulation depth and CT blockade

Across 80 neurons there were significant changes between SAM_Δ100%_ and SAM_Δ25%_ in total spikes in response to both random and repeated trials of stimuli across *f*_mods_, (Table 5). No significant differences in total spikes between SAM_Δ25%_ and SAM_Δ25%_ + CT blockade were noted for randomly presented trials (Table 5). For repeating trials across *f*_mods,_ a switch from SAM_Δ100%_ to SAM_Δ25%_ showed no significant differences in total spikes (731.3 ± 46.3 vs. 693.5 ± 45.1) (Table 5). However, a significant decrease in total spikes was noted when repeating trials across *f*_mod_ were switched from SAM_Δ100%_ to SAM_Δ25%_ to SAM_Δ25%_ + CT blockade (Table 5).

**Table 5:** An average of total spike count and Bonferroni-corrected p-values across all neurons to standard and weakly modulated stimuli presented in random or repeating order.

|  | Total spike count |  |  | p-value* |  |  |
| --- | --- | --- | --- | --- | --- | --- |
| *Presentation order* | $SAM_{\Delta 100\%}$ | $SAM_{\Delta 25\%}$ | $SAM_{\Delta 25\%} + \text{CT blockade}$ | $SAM_{\Delta 100\%} \text{ vs. } SAM_{\Delta 25\%}$ | $SAM_{\Delta 25\%} \text{ vs. } SAM_{\Delta 25\%} + \text{CT blockade}$ | $SAM_{\Delta 100\%} \text{ vs. } SAM_{\Delta 25\%} + \text{CT blockade}$ |
| Random | 839.2±54.6 | 675.6±45.1 | 708.6±50.3 | 0.000005 | 0.549 | 0.001 |
| Repeating | 731.3±46.3 | 693.5±45.1 | 625.8±50.2 | 0.618 | 0.011 | 0.0024 |

Each row represents the average total spike count to $SAM_{\Delta 100\%}$ vs. $SAM_{\Delta 25\%}$ and the
\*Bonferroni-corrected p-values following the repeated measures ANOVA for comparisons. Comparisons were made between $SAM_{\Delta 100\%}$ vs. $SAM_{\Delta 25\%}$ , and $SAM_{\Delta 25\%}$ vs. $SAM_{\Delta 100\%}$ vs. $SAM_{\Delta 100\%}$ vs. $SAM_{\Delta 25\%} + \text{CT blockade}$ , and $SAM_{\Delta 100\%}$ vs. $SAM_{\Delta 25\%} + \text{CT blockade}$ in all the of neurons (n=80).

## Discussion

Previous studies found that both aging and decreased modulation depth, presumptively reducing the salience/fidelity of the ascending temporal code, increased responses to a repeating modulated signal, suggesting engagement of top-down, cognitive and mnemonic resources (Cai *et al*., 2016b; Kommajosyula *et al*., 2019). The present study used optogenetic CT blockade to test whether repetition-enhancement in response to less distinct temporal stimuli was due to the increased involvement of top-down CT resources. In order to maintain speech understanding, older individuals have been shown to increase use of cognitive and memory resources (Bidelman *et al*., 2019a; Roque *et al*., 2019). The impact of aging can be simulated in humans and in animal models by decreasing the temporal clarity of the stimulus. Reducing modulation depth of a SAM stimulus changes the rate and synchrony of the up-stream code introducing temporal jitter (Pichora-Fuller *et al*., 2007; Malone *et al*., 2010; Dimitrijevic *et al*., 2016; Mamo *et al*., 2016). A less temporally distinct ascending acoustic code is thought to engage top-down cognitive resources by generating predictions to support decoding of modulated speech-like signals (Peelle & Wingfield, 2016; Pichora-Fuller *et al*., 2017; Caspary & Llano, 2018; Recanzone, 2018). Consistent with human and animal studies, the present study finds that weakening periodicity cues by decreasing modulation depth (SAM_Δ100%_ to SAM_Δ25%_) decreased the percentage of neurons showing temporal phase-locking to the SAM envelope(Pichora-Fuller *et al*., 2007; Malone *et al*., 2010; Parthasarathy & Bartlett, 2011; Mamo *et al*., 2016; Kommajosyula *et al*., 2019; McClaskey *et al*., 2019). Previously we found that jittering the SAM envelope with a 1.0kHZ centered noise produced similar levels of repetition-enhancement to the SAM_Δ25%_ used in the present study (Kommajosyula *et al*., 2019).) CT blockade did not alter temporal locking of units to the SAM_Δ25%_. The lack of CT blockade changes on temporal locking contrasts to changes observed in SAM rate coding suggesting that CT projections do not play a significant role in temporal coding using this stimulus paradigm (Bartlett & Wang, 2007; Felix *et al*., 2018).

In response to repeating modulated stimuli, decreasing temporal clarity by decreasing modulation depth changed single unit rate responses from adapting to responses showing repetition-enhancement to the repeating modulated SAM stimulus. The switch to increasing responses to less temporally distinct repeating stimuli was blocked/reversed by optical inhibition of CT projections, thought to provide top-down resources to the MGB (Homma *et al*., 2017; Parras *et al*., 2017). A majority of MGB units showed the largest increases in repetition enhancement at higher SAM *f*_mod_ rates (> 128 Hz).

### Temporal distinction and top-down resource usage

The present study used SAM_Δ25%,_ as a surrogate for a diminished acoustic cue that is poorly detected and discriminated in the ascending code in human and animal models of aging (Strouse *et al*., 1998; Nelson & Carney, 2006; Harris & Dubno, 2017). These findings are also consistent with studies modeling aging in young humans with normal hearing and studies of auditory processing of less-distinct stimuli that reveal perceptual deficits due to decrease precision of temporal coding (Shannon *et al*., 1995; Krishna & Semple, 2000; Pichora-Fuller *et al*., 2007; Malone *et al*., 2010; Jorgensen & Dau, 2011; Parthasarathy & Bartlett, 2011; Dimitrijevic *et al*., 2016; Anderson *et al*., 2020; Erb *et al*., 2020).

Previous studies suggest that salience is multidimensional, nonlinear and context-dependent (Kayser *et al*., 2005; Huang & Elhilali, 2017). Based on the context, cortical structures generate predictions of the upcoming sensory stimuli as postulated by predictive coding theory (Mumford, 1992; Koelsch *et al*., 2019). If the prediction and ascending sensory signals do not match, a prediction error should be generated (Auksztulewicz & Friston, 2016). Prediction error is a mechanism to strengthen the internal representation of less temporally distinct stimuli which may lead to generation of a better prediction upon the next repetition (Rao & Ballard, 1999). Studies have suggested increased use of predictive coding in order to cope with less-distinct stimuli or aging accompanied by a less temporally distinct signal to noise ratio (Heinemann *et al*., 2011; Peelle & Wingfield, 2016; Bidelman *et al*., 2019a; Bidelman *et al*., 2019b; Presacco *et al*., 2019; Price *et al*., 2019; Saderi *et al*., 2020). Electrophysiological and fMRI studies suggest a role for repetition suppression/adaptation to repeating stimuli in support of image sharpening and perceptual priming (Gross *et al*., 1967; Dolan *et al*., 1997; James *et al*., 2000; Grill-Spector *et al*., 2006; Näätänen *et al*., 2007). The present findings suggest that for a sensory signal whose features are unclear, adaptation would be counterproductive, whereas repetition-enhancement could potentially facilitate identification of the unclear signal and its characteristics.

The present findings and two prior studies strongly support the idea of CT-mediated transmission of intracortical signals leading to repetition-enhancement (Cai *et al*., 2016b; Kommajosyula *et al*., 2019). Nearly 80% (56/71) of the neurons showed increases in PR, indicating relative increases in unit responses to a repeating stimulus, especially at higher *f*_mods_. MGB units showing the largest repetition enhancement effects (PPI > 0.3) showed increases in firing rates with each successive repeating trial of less-distinct stimuli at higher *f*_mods_ (Fig. 7). SSA studies using short tone-burst stimuli show significantly less adaptation across trials in awake animals, suggesting that top-down projections may reduce SSA in IC and MGB as suggested in the present study and (Antunes *et al*., 2010; Richardson *et al*., 2013; Ayala *et al*., 2015; Duque & Malmierca, 2015; Cai *et al*., 2016a; Yaron *et al*., 2020). The increase in discharge rate with repetition is best explained by a buildup in the strength of the top-down/CT-mediated contribution to the MGB response (Fig. 8B). This is supported by significant decreases in the preference ratios (Figs. 3&5), and trial-by-trial enhancement (Fig. 7) which could be blocked during repeating SAM_Δ25%_ stimuli. The level of adaptation seen with CT blockade during less-distinct stimuli was comparable or greater than seen with the SAM_Δ100%_ stimuli (Figs. 4&5) suggesting blockade of an some on-going level of top-down resource engagement even during a temporally clear SAM_Δ100%_ stimulus. We suggest that CT blockade reduces the ability to convey cortical estimates of the stimulus to MGB neurons, rendering the MGB neurons less sensitive to mismatch/prediction error. (Fig. 8C).

**Fig. 8.**
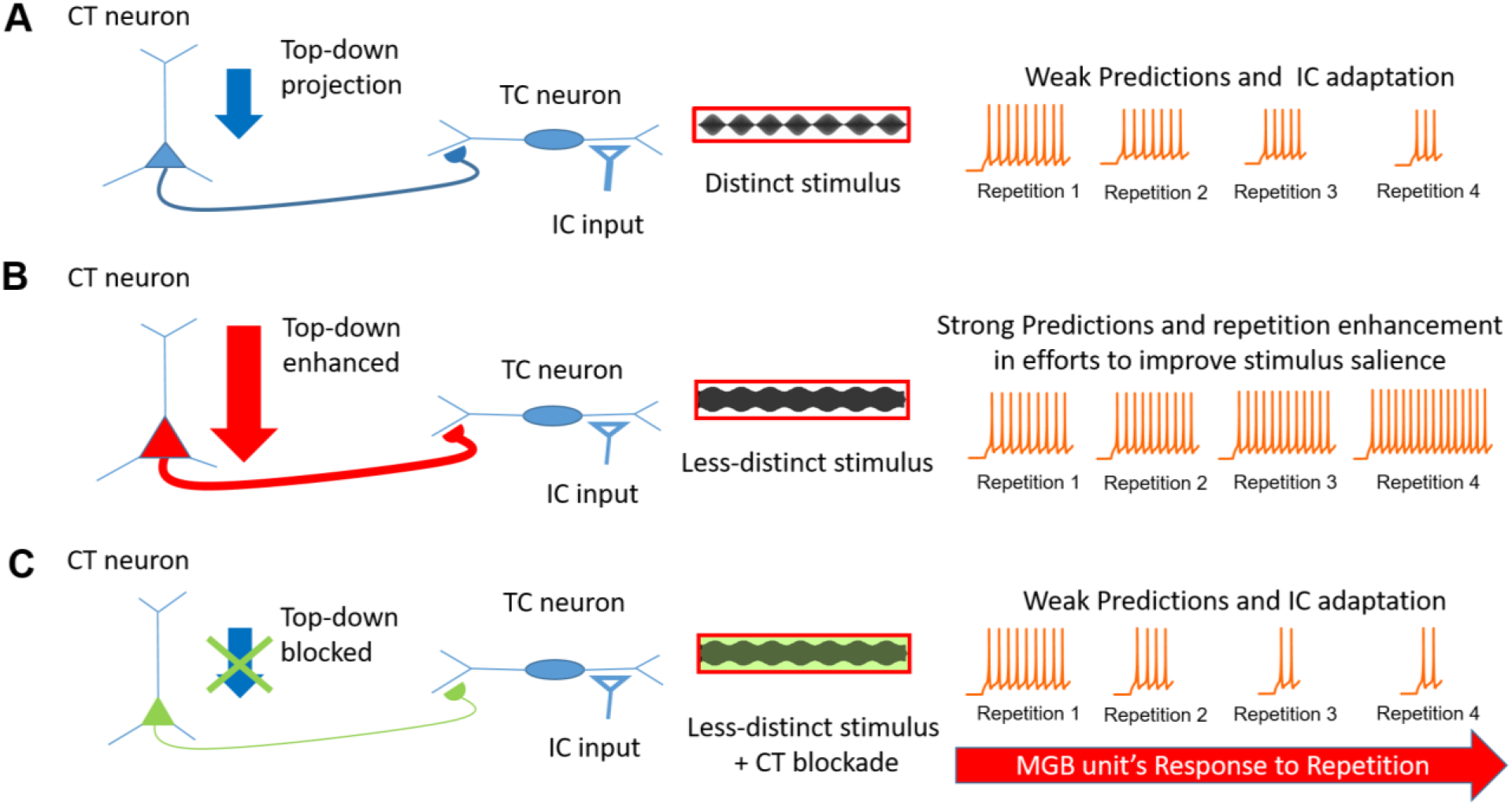
Salience based generation of predication errors in auditory thalamus: An upcoming sensory signal from inferior colliculus (IC) at the level of medial geniculate body (MGB) could interact with a top-down prediction from cortex, and generate prediction error component. The upcoming sensory signals (spikes) generated in response to distinct stimuli, are matched by the top-down predictions and hence little to less generation of prediction error component upon repetition of the distinct stimuli (A). The spike signals to weakly modulated stimuli fail to match the predictions, hence generation of prediction error increases upon repetition until the occurrence of a correct prediction based on the new internal representation formed by feedback from previous prediction error signals. This phenomenon is observed as an increase in response to each repetition (repetition enhancement) (B). CT blockade with weakly modulated stimuli, leads to blockade of delivery of predictions Io MGB, and possibly erroneous prediction error signals and adaptive spike responses (C).

Significant changes in PPI were found in the ventral and dorsal MGB divisions, but not the medial subdivision of the MGB (Fig. 4). The absence of significant changes in the medial subdivision reflect the differential inputs, intrinsic properties and/or connectivity patterns of dorsal MGB neurons, such that they receive different and more widespread CT projections (Smith *et al*., 2007). However, some caution should be exercised in the interpretation of the subdivision findings since recorded neurons were not dye marked and absolute location was only approximated using a template (see methods).

In conclusion, we found that less temporally distinct stimuli increased the preference for repeating modulated signals, i.e. emergence of repetition-enhancement, while blockade of CT projections led to reversal of this effect. In traditional predictive coding theory, an error signal between cortical prediction and incoming sensory inputs generates spiking activity that diminishes as the sensory and prediction templates match, with the mechanisms of this operation not fully understood. The present results are consistent with the idea that a less-distinct acoustic signal leads to the generation of a prediction component similar to what might be seen with phonemic restoration (Bologna *et al*., 2018; Jaekel *et al*., 2018). Cortiothalamic feedback to MGB may serve to amplify weak but predictable features in order to generate a more reliable stimulus template for subsequent predictions, leading to improved detection of changes. We suggest that CT blockade led to a decrease in higher order/top-down information received by MGB neurons, leading to a decrease in corticothalamic mediated repetition-enhancement.

## Author contribution

SPK: Study concept, design, data acquisition, analyses, interpretation, manuscript drafting, and revision; ELB: Data analyses, interpretation and manuscript revision; LL: Confocal imaging, surgical assistance and manuscript revision; RC: Study concept and manuscript revision; DMC: Study concept, design, supervision, data interpretation and manuscript writing and revision. SPK’s present address: Department of Pharmacy, Birla Institute of Technology and Science, Pilani, Hyderabad Campus, Telangana, 500078, India

## Conflict of interest

The authors declare no competing financial interests

## Acknowledgements

This work was supported by National Institute on Deafness and Other Communication Disorders DC000151 to D.M.C. We thank the National Institute on Aging for providing FBN rats; Kevin Brownell for data reduction, Dr. Kristin Delfino for statistical analysis; Dr. Ken Hancock for design and continued development of our stimulus/acquisition system, Lydia Howes for proof reading and Dr. Laurel Carney for suggestions on an earlier version of the manuscript.

## Notes

### Competing Interest Statement

The authors have declared no competing interest.

